# Metabolomic Signatures of Extreme Old Age: Findings from the New England Centenarian Study

**DOI:** 10.1101/2025.09.10.675341

**Authors:** Stefano Monti, Michael S. Lustgarten, Ziwei Huang, Zeyuan Song, Dylan Ellis, Qu Tian, Luigi Ferrucci, Noa Rappaport, Stacy L. Andersen, Thomas P. Perls, Paola Sebastiani

## Abstract

The New England Centenarian Study (NECS) provides a unique resource for the study of extreme human longevity (EL). To gain insight into biological pathways related to EL, chronological age and survival, we used an untargeted serum metabolomic approach (> 1,400 metabolites) in 213 NECS participants, followed by integration of our findings with metabolomic data from four additional studies. Compared to their offspring and matched controls, EL individuals exhibited a distinct metabolic profile characterized by higher levels of primary and secondary bile acids – most notably chenodeoxycholic acid (CDCA) and lithocholic acid (LCA) – higher levels of biliverdin and bilirubin, and stable levels of selected steroids. Notably, elevated levels of both bile acids and steroids were associated with lower mortality. Several metabolites associated with age and survival were inversely associated with metabolite ratios related to NAD+ production and/or levels (tryptophan/kynurenine, cortisone/cortisol), gut bacterial metabolism (ergothioneine/ trimethylamine N-oxide, aspartate/quinolinate), and oxidative stress (methionine/methionine sulfoxide), implicating these pathways in aging and/or longevity. We further developed a metabolomic clock predictive of biological age, with age deviations significantly associated with mortality risk. Key metabolites predictive of biological aging, such as taurine and citrate, were not captured by traditional age analyses, pointing to their potential role as biomarkers for healthy aging. These results highlight metabolic pathways that may be targeted to promote metabolic resilience and healthy aging.

## INTRODUCTION

Aging is characterized by complex and dynamic molecular changes that affect nearly every physiological system and eventually emerge as chronic diseases^1–5^. Although these changes occur in the majority of individuals, many centenarians avoid or substantially delay the onset of major age-related diseases^6–9^. These individuals provide a powerful model for studying the biological underpinnings of healthy aging and extreme longevity, offering insights into how to increase *healthspan*, and how to delay or compress the period of life burdened by age-related diseases^10,11^.

Metabolomics, the comprehensive analysis of small-molecule metabolites in biological samples, provides unique insights into physiological and pathological processes that occur with age^12,13^. As metabolites represent the end products of gene-environment interactions, they are especially informative to understand the integrated biochemical state of an organism. Numerous metabolomics studies reported changes in lipid metabolism and amino acid metabolism, as well impaired mitochondrial health and inflammation ^14–16^. However, relatively few have focused specifically on extreme longevity, which is here defined as living beyond the 99^th^ percentile of a subject’s birth cohort^17^.

Studies of serum metabolomics have reported altered levels of sphingolipids, acylcarnitines, and amino acids in centenarians compared to younger or elderly controls, suggesting roles for mitochondrial efficiency, lipid remodeling, and reduced pro-inflammatory signaling in longevity. For example, Montoliu *et al*. (2014)^18^ and Collino *et al.* (2013)^19^ identified metabolite signatures linked to healthy aging and long-lived phenotypes in European cohorts, while recent work in Japanese centenarians highlighted potential roles for gut microbiota-derived metabolites and anti-inflammatory bile acids^20,21^. A recent study by Sebastiani *et al.* (2024)^22^ used untargeted metabolomics in the Long-Life Family Study (LLFS) to define metabolite signatures of chronological age, aging, and survival, identifying changes in amino acids, nucleotides, lipids, and markers of mitochondrial function, and provides an important foundation for understanding systemic metabolic change in a large cohort enriched for long-lived individuals.

Despite these advances, major gaps remain: most studies assay a relatively limited number of metabolites, lack replication across populations, or lack survival follow-up. Toward addressing these gaps, we conducted an untargeted serum metabolomic study of 213 New England Centenarian Study (NECS) participants^17^. A subset of 408 metabolites from these data were previously analyzed in a metabolomic study of 2,764 LLFS participants^22^. In this study, we extended our analysis to the larger set of 1,495 metabolites that were measured in the NECS cohort. Using deep untargeted metabolomic profiling and integrating our results with those from independent metabolomic datasets, we aimed to: (1) define metabolomic signatures associated with exceptional longevity (EL); (2) evaluate whether these signatures are conserved across independent cohorts; (3) distinguish age-associated from EL-associated metabolic signatures by contrasting results from studies enriched for EL individuals and studies excluding EL subjects, and 4) through triangulation with survival, identify those metabolites more likely to contribute to EL. Finally, we built a metabolomic clock estimating biological age, to assess its ability to capture deviations from chronological age predictive of survival. By analyzing and integrating cross-sectional data, this study seeks to uncover molecular pathways underlying exceptional longevity, and to identify metabolite signatures that may inform future strategies to promote healthy aging and healthspan extension.

## RESULTS

### Serum Metabolomic Analysis of a Population Uniquely Enriched for EL Subjects

We analyzed 1,495 metabolites (Table S1) in 213 NECS participants^17^, including 70 EL subjects (centenarians, see M&M), 80 offspring, and 63 controls (spouses of centenarian offspring or members of families without longevity), with their demographics summarized in Table 1. We integrated our results with previously published metabolomic data from two cohorts that included long-lived individuals – LLFS (*n* = 2,764)^22,23^ and Xu *et al.* (*n* = 382)^21^, and two cohorts that did not include EL individuals - Arivale (*n* = 634)^24^ and the Baltimore Longitudinal Study of Aging (BLSA; *n* = 1,135)^25^. These datasets were generated using different metabolomic platforms^21–25^, each with distinct coverage of metabolites and age distributions (Table S2). The combined datasets encompassed 5,128 individuals across diverse age ranges (Figure 1A), with a comparison of age distributions between NECS and the other cohorts shown in Figure 1B. Principal component analysis (PCA) of the NECS samples revealed a clear separation of centenarians from the rest of the cohort (Figure 1C), suggesting a distinct metabolomic profile.

**Figure 1.**
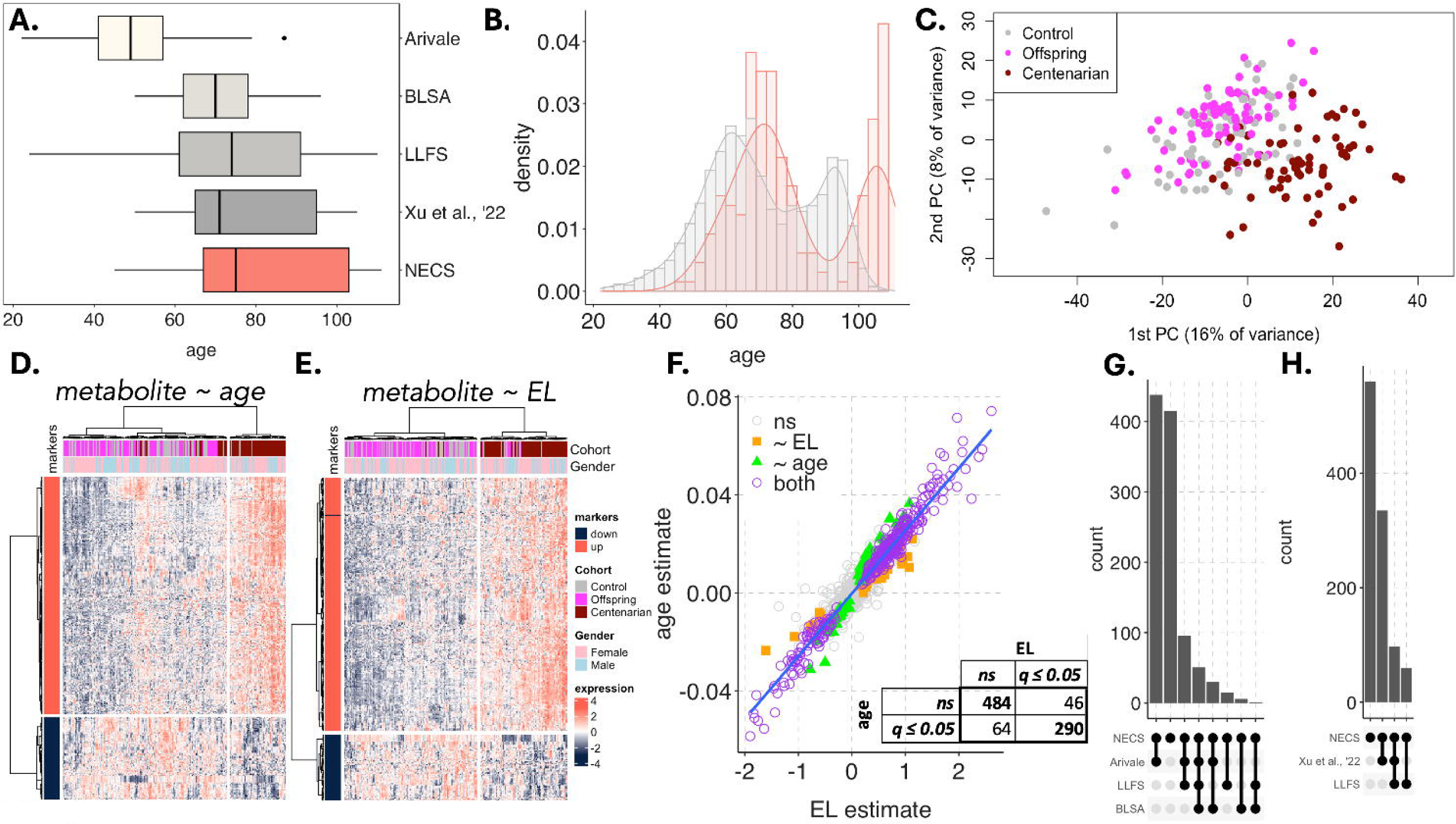
Summary statistics of the datasets analyzed and signature identification. **A)** Age distribution within each dataset. **B)** Comparison of age distributions in NECS vs. the other datasets. **C)** NECS metabolomics projection onto the first two principal components. **D,E)** Heatmaps of the significant (q ≤ 0.01) age and EL markers, respectively. **E)** Comparison of the metabolite estimates (beta coefficients) from the age and EL-based analyses, and confusion matrix of significant markers in the two analyses (ns = non-significant). **G)** Significant age-associated and EL-associated metabolite overlap among the datasets compared.

**Table 1:** NECS subjects’ demographics.

|  | Cohort |  |  |
| --- | --- | --- | --- |
| Characteristics | Control N=63 (30%) | Offspring N=80 (37%) | Centenarian N=70 (33%) |
| Age at Blood Draw | 71 ± 7.9 (52, 88) | 71 ± 9.2 (45, 90) | 105 ± 3.8 (90,111) |
| Age at Last Follow-up | 81 ± 7.1 (64, 100) | 83 ± 8.7 (56, 100) | 108 ± 3.4 (96, 115) |
| Years of Education | 16.1 ± 2.9 (10, 22) | 15.6 ± 3.0 (12, 25) | 11.6 ± 4.1 (2, 21.5) |
| Gender |  |  |  |
| Male | 34 (54%) | 53 (66%) | 47 (64%) |
| Female | 29 (46%) | 27 (34%) | 23 (33%) |

### Metabolomic Signature of Age Recapitulates and Expands upon Previous Findings

We identified 360 metabolites that were significantly associated with chronological age (q-value ≤ 0.05), of which 263 were positively and 93 were negatively associated with age (Figure 1D, Table S3).

Metabolite-set (MSet) over-representation analysis (Tables S4 and S5) identified sixteen metabolites within the ***Urea Cycle and Related Metabolites*** category that significantly increased with age, including citrulline, proline, trans-4-hydroxyproline, dimethylarginine, homocitrulline, N-acetylcitrulline, pro-hydroxy-pro, 2-oxoarginine, and others. These are linked to processes such as endothelial dysfunction, oxidative stress, mitochondrial impairment, and aging-related inflammation^14,26–28^. Included in this category were several *Methylation and Betaine Derivatives,* including N-methylproline, N-methylhydroxyproline, and N,N,N-trimethyl-alanylproline betaine, which also increased with age, as did 3-amino-2-piperidone. In agreement with previous studies^29–31^, ***Tyrosine Metabolism*** metabolites, including homovanillic acid, vanillylmandelate, and other tyrosine derivatives (e.g., N-formylphenylalanine, 1-carboxyethyltyrosine, 3-HPLA, p-cresol sulfate) increased with age, many of which are uremic toxins. Fifteen metabolites, classified as ***Dicarboxylic Acids***, including methylmalonate, suberate, and azelate, increased with age, reflecting mitochondrial dysfunction and renal function decline^32–35^. Eight ***Sugar Acid*** metabolites, including glucuronate and 2R,3R-dihydroxybutyrate, increased with age, implicating oxidative stress and impaired energy metabolism^36,37^. Eleven ***Long Chain Polyunsaturated Fatty Acids (PUFAs***), seven ***Fatty Acid Metabolism (Acyl Choline)*** metabolites, and six ***Long Chain Monounsaturated Fatty Acids (MUFAs)*** declined with age.

*<u>Replication</u>*. Of the 888 metabolites with known identity, 577 were measured in at least one of the other datasets – Arivale, LLFS, and BLSA – and within this set, of the 220 metabolites significantly associated with age in NECS, 111 were also significantly associated with age in the same direction in at least one other study (91 and 20 positively (+) and negatively (-) associated with age, respectively), with 22 metabolites (20+ and 3-) replicated in two studies, including choline (increasing) and pregnenediol sulfate and pregnenolone sulfate (decreasing), and with four metabolites (3+ and 1-) replicated in all three studies, including tryptophan, which consistently declined during aging (Figure 1G, Figure S1A, Table S3).

#### EL Signature

We compared the metabolomic profiles of EL participants versus offspring and controls. This analysis yielded 336 metabolites that were significantly associated with EL (q-value ≤ 0.05), of which 257 (79) were more (less) abundant in the EL group than in controls (Figure 1E, Table S3). Comparison of the age and EL-signatures identified 290 metabolites that were shared by both profiles, whereas 46 and 64 unique metabolites were significantly associated only with the EL and the aging signatures, respectively (Figure 1F). Concordantly, MSet enrichment analysis (Tables S4 and S5) identified many of the same categories enriched in the age signature (Figure 3).

*<u>Replication</u>*. Of the 888 metabolites with known identity measured in NECS, 505 were measured in at least one of the other datasets that included EL subjects – Xu *et al.*^21^, and LLFS – and within this set, of the 211 metabolites significantly associated with EL in NECS, 182 were also significantly associated in the same direction with EL in at least another study (148 and 34 positively and negatively associated with EL, respectively), and 25 (21+ and 4-) were replicated in two studies (Figure 1H, Figure S1B, Table S3).

#### Module-Based Analysis

Given the large number of metabolites significantly associated with the studied phenotypes, and that many of these metabolites are highly correlated, we next identified and annotated *modules* of highly correlated metabolites and calculated for each module an overall score. Then, we tested whether these scores were associated with age-related phenotypes.

#### Module Identification and Annotation

We applied WGCNA to identify modules of co-expressed metabolites, which yielded 31 modules ranging in size from 11 to 133 metabolites (Figure S2). Module composition, and their annotation based on over-representation analysis using the Metabolon- and Refmet-based MSets (see M&M), are summarized in Table S6. For example, the “darkred” module (size = 16) was highly enriched for ***Secondary*** (q < 1.5E-13) and ***Primary*** (q < 1.0E-5) ***Bile Acids***, while the “pink” module (size = 35) was highly enriched for ***Androgenic*** (q < 1.0E-15)*, **Pregnenolone*** (1.0E-9)*, and **Progestin*** (q < 0.0002) ***Steroids***. Altogether, 30 out of 31 modules were significantly enriched for at least one Metabolon or RefMet MSet, with minimal overlap in terms of the modules’ top enrichment, thus supporting their utilization for downstream analysis.

#### Module-Based Age and EL markers

Sample-specific module scores were computed using GSVA^38^, yielding a 31-by-213 matrix of module-by-sample scores to be used for regression and differential analysis, based on the same linear models and covariates that were used for the metabolite-level analyses. Age and EL analyses yielded 14 (6+ and 8-) and 14 (6+ and 8-) significantly associated modules, respectively, and the results are summarized side-by-side in Figure 2, with the modules annotated based on their top Metabolon MSet enrichment (Table S7). ***Tyrosine Metabolism***, ***Leucine, Isoleucine, and Valine Metabolism*,** and ***Secondary Bile Acids*** were the top three modules positively associated with EL; ***Hemoglobin and Porphyrin Metabolism***, ***Fatty Acid, Amino***, and ***Androgenic Steroids*** were the top three modules negatively associated with EL. Analogous analyses were repeated based on manually curated “modules” corresponding to the Metabolon and RefMet MSets (see M&M), obtaining similar results (Figure S3, Tables S8 and S9).

**Figure 2.**
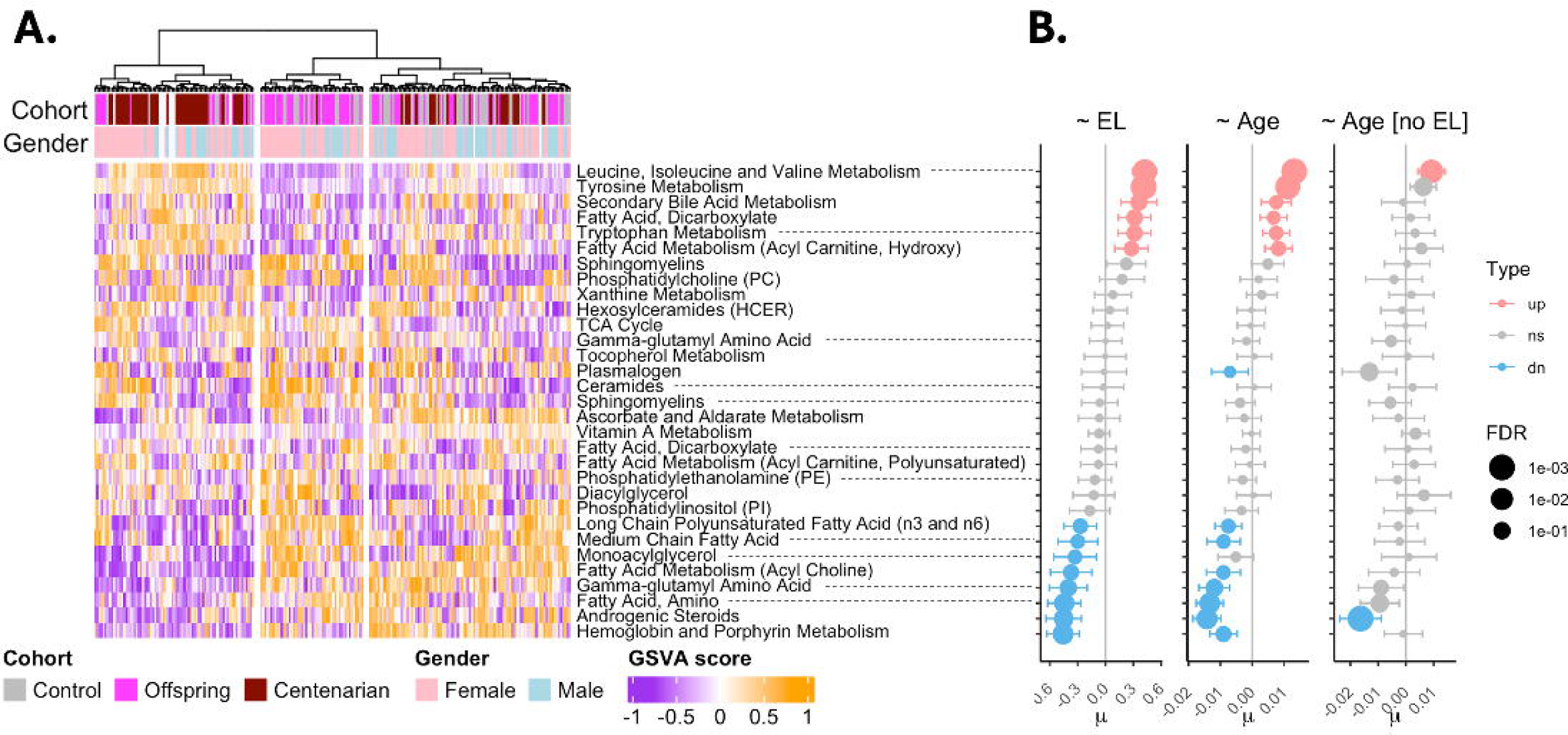
Module-based analysis. **A)** Heatmap of module scores. **B)** Module-based reegression on EL status and age (with and w/o EL subjects included). Modules (y-axis) derived by WGCNA, and sample-level module scores computed using GSVA. Each module (y-axis) is annotated based on the top enrichment of the Metabolon MSets.

The module-based age regression was then repeated on the subset of NECS samples after excluding the EL subjects. Several of the modules significantly associated with EL were not associated with age in this analysis (Figure 2. See Figure S1D for a comparison of the effect sizes). However, the limited sample size (n = 133), and the fact that more than half of the non-EL subjects were EL offspring prevented us from drawing strong conclusions about the distinction between age and EL signals, a point to which we now turn.

### Disentangling Age and EL Metabolomic Signatures

To distinguish between age and EL signals, we integrated our findings with published metabolomic studies of aging that did not include EL subjects (non-EL studies, for short). This approach leveraged metabolite measurements common to both studies and identified metabolites significantly associated with EL exclusively in our dataset, or exclusively with age in non-EL studies, thereby highlighting them as potential extreme longevity (*EL-only*) and aging (*age-only*) markers, respectively.

#### Age-only Markers

To identify age-only markers, we compared our EL signature to age markers from datasets that did not include EL subjects (non-EL studies: Arivale and BLSA; see M&M section). This yielded a set of 78 identified metabolites, of which 60 (18) were positively (negatively) associated with age (Table S3), with the top four markers including *alpha-tocopherol*, *eicosenoylcarnitine (c20:1)\**, and *sphingomyelin (d17:2/16:0, d18:2/15:0)\** (positively associated with age), and *16a-hydroxy dhea 3-sulfate* (negatively associated with age). Selected age-only metabolites are shown in Figure 4a,b. Of note, while tryptophan did not meet our stringent inclusion criteria, its increased abundance was significantly associated with age in non-EL studies but was not significant (q = 0.07) in the NECS. MSet-based over-representation analysis identified several classes of metabolites that were significantly (and exclusively) associated with age, including negative associations for multiple ***Progestin, Androgenic,*** and ***Pregnenolone Steroids***, as were multiple ***Leucine, Isoleucine and Valine*** metabolites (q < 0.0004). Conversely, many metabolites in the ***Acyl Carnitine Mono-*** and ***Poly-unsaturated*** and ***Long Chain Saturated*** groupings were positively associated with age (Figure 3A, Tables S4 and S5). Table 2 reports the age-only markers with q≤0.01 (see Table S3 for the full list of q≤0.05 markers).

**Figure 3.**
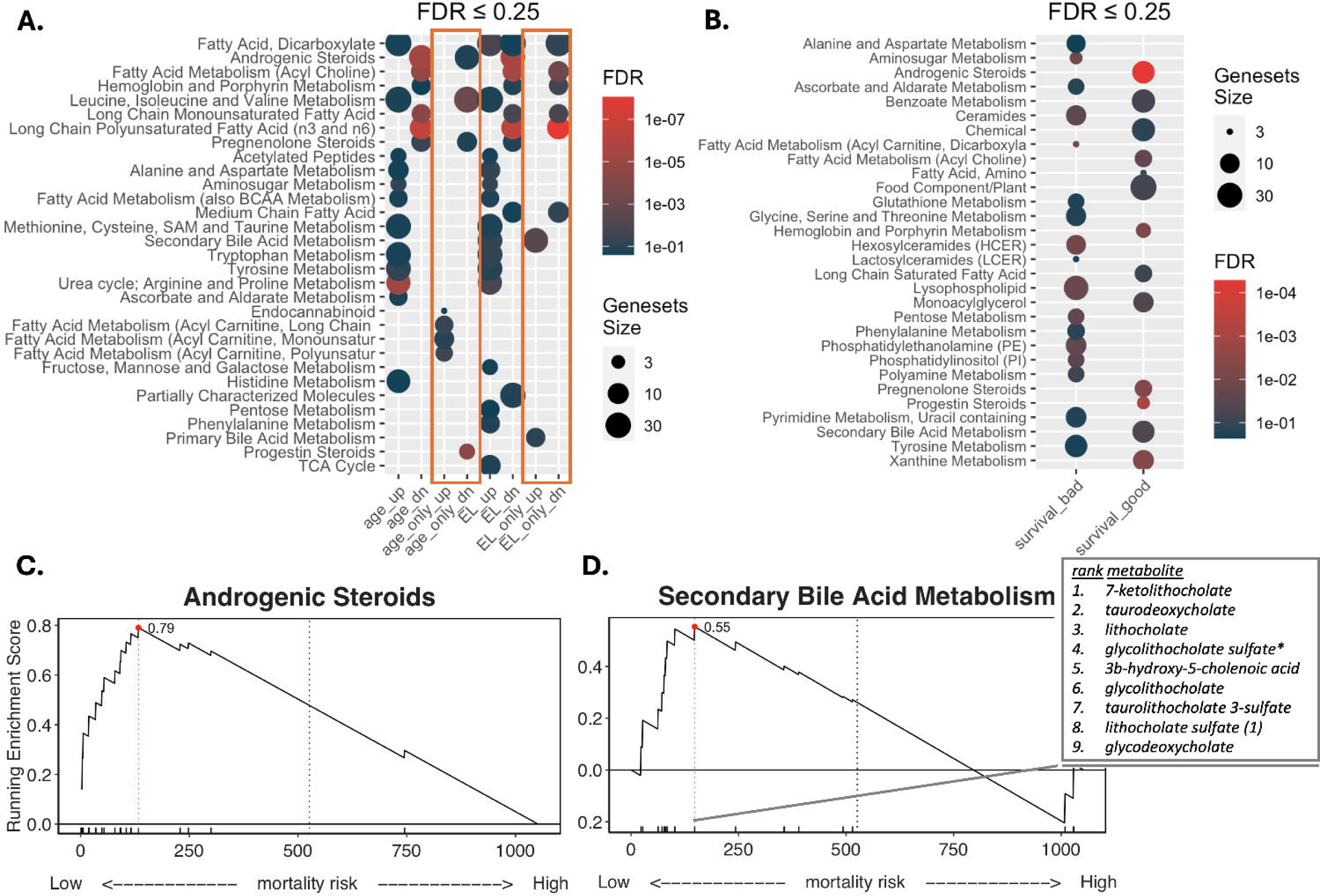
Metabolite signatures annotation, using Metabolon-derived ‘SUB_PATHWAY’ classes, by. **A)** Over-representation analysis; and **B)** rank-based Kolmogorov-Smirnov test. **C)** Rank-based enrichment plot for the Androgenic Steroids MSet. **D)** Rank-based enrichment plot for the Secondary Bile Acids MSet, with the leading-edge metabolites listed.

**Figure 4.**
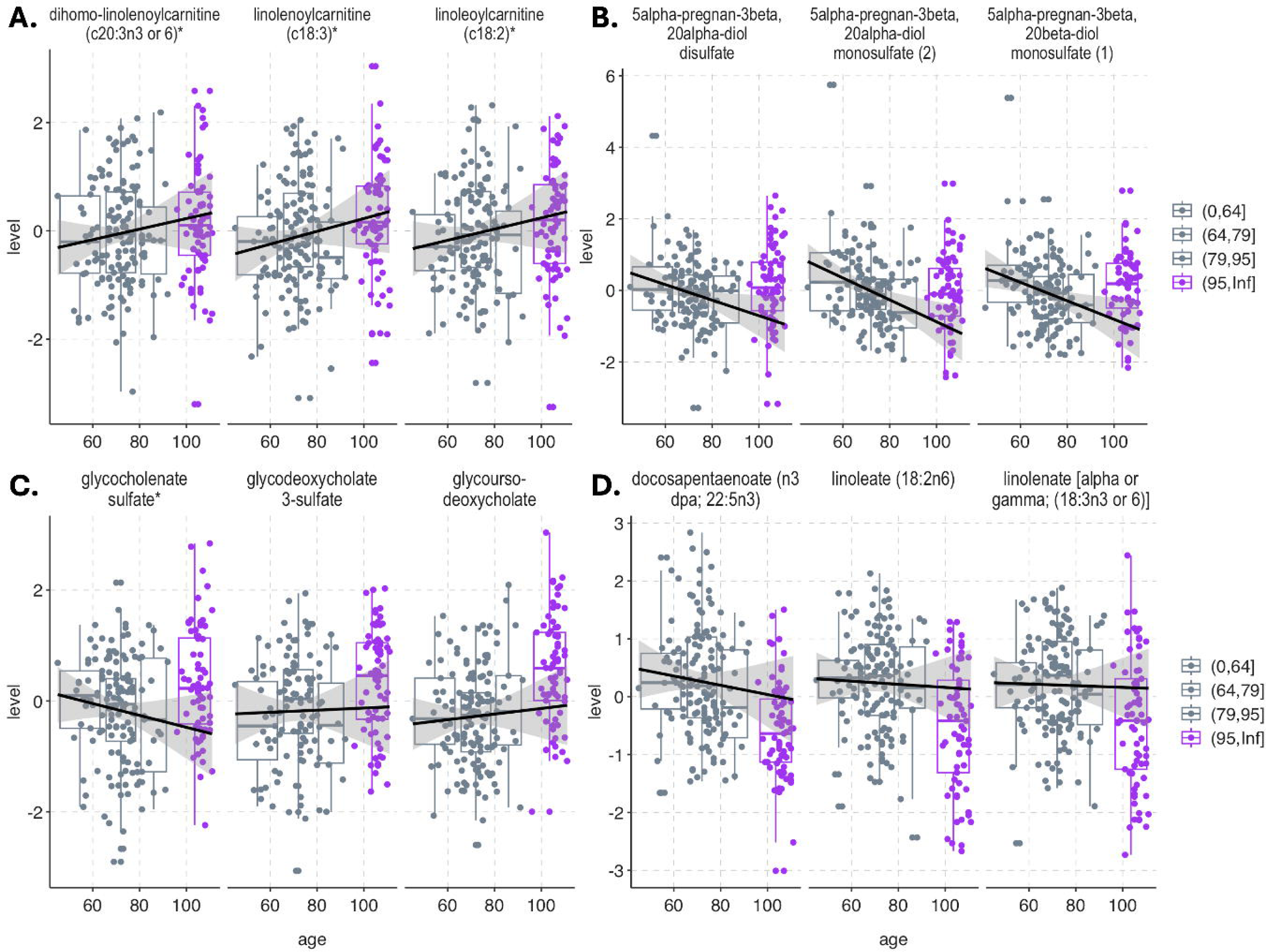
Disentangling age and EL signals. **A-B)** Selected “Age-only” (i.e., “bucking the trend”) metabolites increasing and decreasing with age, respectively. **C-D)** Selected “EL-only” metabolites more and less abundant in EL subjects compared to non-EL subjects, respectively. Trend lines were fitted by excluding the oldest group. Caveat of visualization: non-EL portion of plots does not fully reflect the statistics used (drawn from non-EL studies, i.e., Arivale and BLSA).

**Table 2:** Age-only markers. List of metabolites exclusively associated with age in non-EL individuals (q≤0.01).

| CHEMICAL_NAME | age.beta | age.q.value | SUPER_PATHWAY | SUB_PATHWAY |
| --- | --- | --- | --- | --- |
| trans-urocanate | -0.00624 | 0.00446 | Amino Acid | Histidine Metabolism |
| 2-hydroxy-3-methylvalerate | -0.00891 | 0.000000242 |  | Leucine, Isoleucine and Valine Metabolism |
| alpha-hydroxyisocaproate | -0.00667 | 0.000000352 |  |  |
| 3-methyl-2-oxovalerate | -0.00457 | 0.00000479 |  |  |
| 4-methyl-2-oxopentanoate | -0.00414 | 0.0000459 |  |  |
| leucine | -0.00177 | 0.000315 |  |  |
| n,n,n-trimethyl-5-aminovalerate | 0.00547 | 0.00159 |  | Lysine Metabolism |
| cysteine | 0.00404 | 0.000728 |  | Methionine, Cysteine, SAM and Taurine Metabolism |
| tyrosine | 0.00282 | 0.0019 |  | Tyrosine Metabolism |
| n-acetylarginine | 0.00716 | 0.0000129 |  | Urea cycle; Arginine and Proline Metabolism |
| trigonelline (N<sup>1</sup>-methylnicotinate) | 0.0163 | 0.0000142 | Cofactors and Vitamins | Nicotinate and Nicotinamide Metabolism |
| N1-methyl-2-pyridone-5-carboxamide | 0.00535 | 0.00213 |  |  |
| alpha-tocopherol | 0.00651 | 3.82E-17 |  | Tocopherol Metabolism |
| pyridoxate | 0.00654 | 0.00235 |  | Vitamin B6 Metabolism |
| 16a-hydroxydhea-3-sulfate | -0.0235 | 1.02E-14 | Lipid | **Androgenic Steroids** |
| androsteroid monosulfate c19h28o6s (1)* | -0.0183 | 9.34E-10 |  |  |
| etiocholanolone glucuronide | -0.0105 | 0.0000791 |  |  |
| ceramide (d18:1/14:0, d16:1/16:0)* | 0.00711 | 0.00000565 |  | Ceramides |
| ceramide (d18:1/20:0, d16:1/22:0, d20:1/18:0)* | 0.00637 | 0.0000142 |  |  |
| ceramide (d18:2/24:1, d18:1/24:2)* | 0.00506 | 0.000235 |  |  |
| cortolone glucuronide (1) | 0.00536 | 0.000409 |  | Corticosteroids |
| palmitoyl dihydrosphingomyelin (d18:0/16:0)* | 0.00389 | 0.0000491 |  | Dihydrosphingomyelins |
| myristoyl dihydrosphingomyelin (d18:0/14:0)* | 0.00378 | 0.00635 |  |  |
| oleoyl ethanolamide | 0.00981 | 0.000000382 |  | Endocannabinoid |
| palmitoyl ethanolamide | 0.00498 | 0.00007 |  |  |
| stearoylcarnitine (c18) | 0.00976 | 8.75E-13 |  | Fatty Acid Metabolism (Acyl Carnitine, Long Chain Saturated) |
| myristoylcarnitine (c14) | 0.00967 | 5.93E-09 |  |  |
| palmitoylcarnitine (c16) | 0.0057 | 0.00000102 |  |  |
| margaroylcarnitine (c17)* | 0.00583 | 0.000151 |  |  |
| eicosenoylcarnitine (c20:1)* | 0.0125 | 2.24E-15 |  | Fatty Acid Metabolism (Acyl Carnitine, Monounsaturated) |
| palmitoleoylcarnitine (c16:1)* | 0.0121 | 2.75E-13 |  |  |
| oleoylcarnitine (c18:1) | 0.00938 | 8.91E-12 |  |  |
| ximenoylcarnitine (c26:1)* | 0.00707 | 0.000000532 |  |  |
| linolenoylcarnitine (c18:3)* | 0.00999 | 1.18E-08 |  | Fatty Acid Metabolism (Acyl Carnitine, Polyunsaturated) |
| linoleoylcarnitine (c18:2)* | 0.00722 | 0.00000228 |  |  |
| arachidonoylcarnitine (c20:4) | 0.00667 | 0.000311 |  |  |
| dihomo-linolenoylcarnitine (c20:3n3 or 6)* | 0.00518 | 0.00604 |  |  |
| butyrylcarnitine (c4) | 0.00573 | 0.0063 |  | Fatty Acid Metabolism (also BCAA Metabolism) |
| n-acetyl-2-aminooctanoate* | 0.012 | 0.000000135 |  | Fatty Acid, Amino |
| 3-carboxy-4-methyl-5-pentyl-2-furanpropionate (3-cmpfp)** | 0.00789 | 0.00000236 |  | Fatty Acid, Dicarboxylate |
| branched chain 14:0 dicarboxylic acid** | 0.0144 | 0.000169 |  |  |
| glycosyl-n-tricosanoyl-sphingadienine (d18:2/23:0)* | 0.003 | 0.000905 |  | Hexosylceramides (HCEr) |
| glycosyl-n-behenoyl-sphingadienine (d18:2/22:0)* | 0.0053 | 0.00105 |  |  |
| 1-linoleoylglycerol (18:2) | 0.00861 | 0.000206 |  | Monoacylglycerol |
| 1-stearoyl-2-oleoyl-gpc (18:0/18:1) | 0.00506 | 1.97E-08 |  | Phosphatidylcholine (PC) |
| 1-stearoyl-2-docosahexaenoyl-gpc (18:0/22:6) | 0.00584 | 0.00000335 |  |  |
| 1-oleoyl-2-docosahexaenoyl-gpc (18:1/22:6)* | 0.00426 | 0.0000356 |  |  |
| pregnenediol disulfate (c21h34o8s2)* | -0.017 | 2.83E-12 |  | **Pregnenolone Steroids** |
| pregnenetriol disulfate* | -0.016 | 7.53E-11 |  |  |
| 5alpha-pregnan-3beta,20beta-diol monosulfate (1) | -0.0298 | 1.57E-11 |  | **Progestin Steroids** |
| 5alpha-pregnan-3beta,20alpha-diol disulfate | -0.024 | 0.000000087 |  |  |
| pregnanediol-3-glucuronide | -0.0196 | 0.0000017 |  |  |
| 5alpha-pregnan-3beta,20alpha-diol monosulfate (2) | -0.0221 | 0.00000719 |  |  |
| sphingomyelin (d17:2/16:0, d18:2/15:0)* | 0.0112 | 5.84E-15 |  | Sphingomyelins |
| sphingomyelin (d18:2/21:0, d16:2/23:0)* | 0.00837 | 4.7E-12 |  |  |
| sphingomyelin (d18:2/14:0, d18:1/14:1)* | 0.00733 | 3.49E-08 |  |  |
| sphingomyelin (d18:1/19:0, d19:1/18:0)* | 0.00625 | 0.00000236 |  |  |
| sphingomyelin (d18:1/18:1, d18:2/18:0) | 0.00275 | 0.0022 |  |  |
| 3-aminoisobutyrate | 0.00517 | 0.00735 | Nucleotide | Pyrimidine Metabolism, Thymine containing |
| o-cresol sulfate | 0.0102 | 1.1E-10 | Xenobiotics | Benzoate Metabolism |
| 3-hydroxyhippurate | 0.0167 | 0.000501 |  |  |
| sulfate* | 0.00251 | 0.000856 |  | Chemical |
| 4-allylphenol sulfate | 0.0158 | 0.000409 |  | Food Component/Plant |
| 1,7-dimethylurate | 0.0144 | 0.00285 |  | Xanthine Metabolism |
| 5-acetyl-amino-6-amino-3-methyluracil | 0.0134 | 0.00608 |  |  |

#### EL-only Markers

The *EL-only signature* was defined as the set of metabolites that were significant in NECS but not significantly associated with age in all non-EL studies in which they were assayed (see M&M). This yielded a set of 67 identified metabolites, of which 41 (26) were more (less) abundant in EL than in non-EL subjects, with 44 (21+ and 24-) metabolites replicated in at least one other study (Table 3, q ≤ 0.01, and Table S3, q ≤ 0.05), and with the top four markers being *n-acetylneuraminate*, *glycochenodeoxycholate*, and *orotidine* (positively associated with EL), and *biliverdin* (negatively associated). Selected EL-only metabolites are shown in Figure 4c,d. Over-representation enrichment analysis identified ***Primary* and *Secondary Bile Acids*** as positively associated, whereas several classes were negatively associated with EL, including ***Long Chain Polyunsaturated Fatty Acids* (n3 and n6)**, ***Long Chain Monounsaturated Fatty Acids***, ***Fatty Acid (Acylcholines)***, and ***Hemoglobin and Porphyrin* metabolites**, among others (Figure 3A, Tables S4 and S5).

**Table 3:**
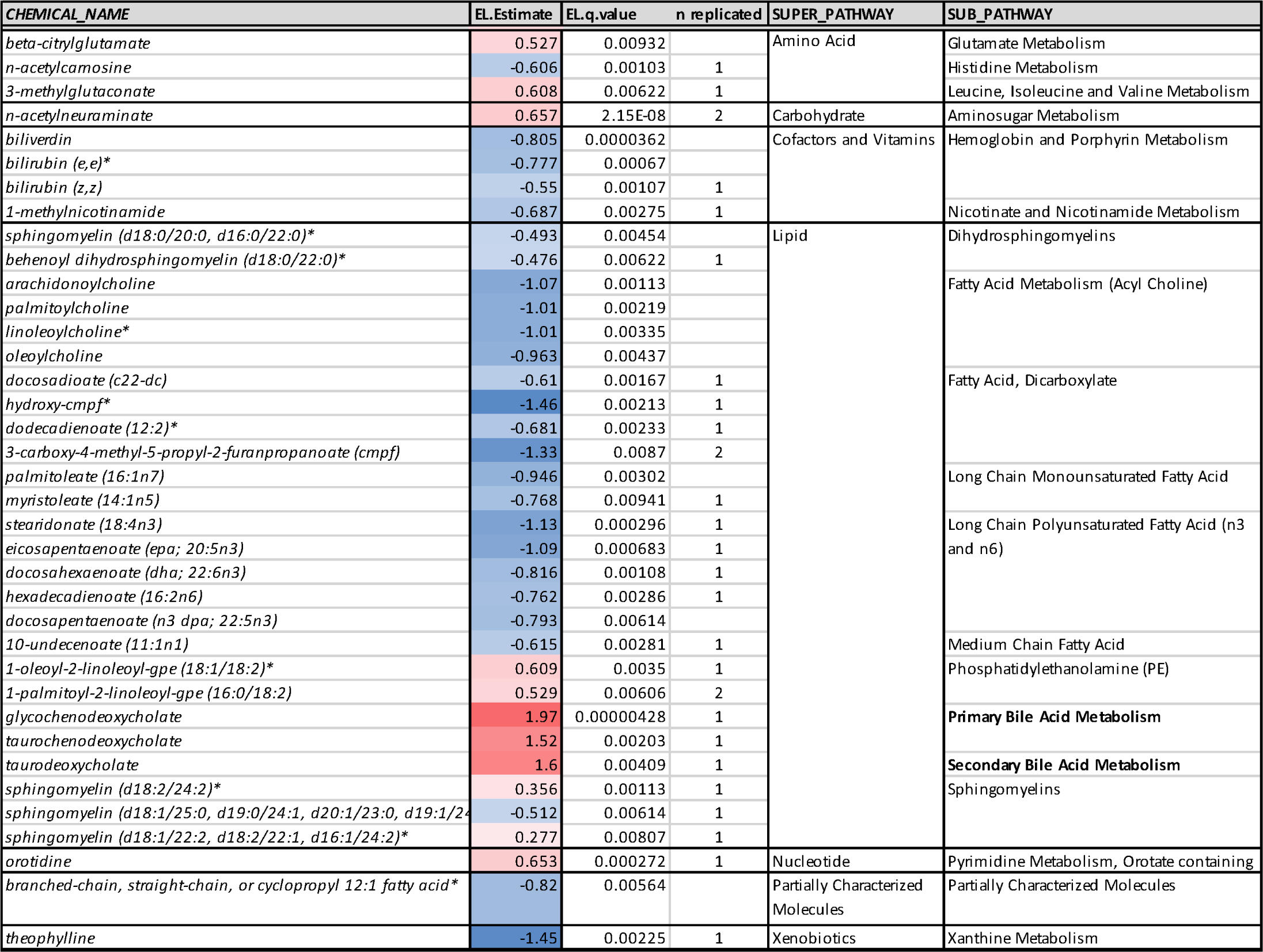
EL-only markers. List of metabolites exclusively associated with EL status (q≤0.01).

#### Prognostic Markers

The availability of follow-up information for NECS subjects allowed us to test for the association of metabolite levels with length of survival past blood collection. To this end, we performed survival analysis based on a Cox proportional hazard model regressing length of follow-up on each metabolite, while controlling for age, gender, education, batch, and serum sample storage duration. None of the individual metabolites reached statistical significance at the multiple hypothesis testing (MHT)-corrected q = 0.05. However, several *classes* of metabolites when tested as a group were significantly associated with mortality risk, based on a Kolmogorov-Smirnov (KS) rank test: higher levels of ***Androgenic, Progestin,*** and ***Pregnenolone Steroids, Secondary Bile Acids, Hemoglobin and Porphyrin Metabolites, Fatty Acids (Acylcholines),*** and ***Monoacylglycerols*** were associated with better survival, whereas higher levels of ***Fatty Acid (Acyl Carnitine, Dicarboxylate) metabolites***, among others, were significantly associated with worse survival (Figure 4B, Tables S10 and S11).

#### Signature Triangulation

To distinguish candidate metabolites contributing to EL from candidate end-of-life (EoL) markers, we triangulated age-only and EL-only signatures with survival signatures. The underlying hypothesis was that metabolites uniquely altered in EL subjects and predictive of reduced mortality risk beyond blood collection are more likely to contribute to EL, whereas those predictive of increased mortality risk are more likely to be EoL markers.

Table 4 summarizes the results of this triangulation. For example, ***Steroids***, especially progestin steroids (q <1.0E-4), were negatively associated with age only in non-EL subjects (i.e., less abundant in older people but not in EL subjects), and ***Bile acids*** were positively associated with EL status only (i.e., more abundant in EL subjects but not in older non-EL subjects). Higher abundances of both metabolite classes were significantly associated with lower mortality risk, indicating their unique expression patterns in EL are linked to better prognosis. Conversely, higher levels of ***Fatty Acids (Acyl cholines),*** and ***Hemoglobin and Porphyrin metabolites*** were also associated with lower mortality risk but were found to be less abundant exclusively in EL subjects (EL-only “down” markers), suggesting these metabolite changes in EL subjects correspond to worse prognosis.

**Table 4:** Signature Triangulation: Combining the significant enrichments from the age- and EL-associated signatures with the significant enrichments from the survival signatures and their directions, we can distinguish candidate EL-associated metabolic processes from candidate End of Life (EoL) processes.

| Super Classes | Classes | Age-Only |  | EL-only |  | Survival |  | EL vs. EoL? |
| --- | --- | --- | --- | --- | --- | --- | --- | --- |
|  |  | Up | Down | Up | Down | Better | Worse |  |
| **Steroid** | Progestin Steroids |  | ✓ |  |  | ✓ |  | EL? |
|  | Androgenic Steroids |  | ✓ |  |  | ✓ |  | EL? |
|  | Pregnenolone Steroids |  | ✓ |  |  | ✓ |  | EL? |
| **Bile Acids** | Secondary Bile Acids |  |  | ✓ |  | ✓ |  | EL? |
|  | Primary Bile Acids |  |  | ✓ |  |  |  | EL? |
| **Lipids** | Fatty Acid (acyl choline) |  |  |  | ✓ | ✓ |  | EoL? |
| **Cofactors & Vitamins** | Hemoglobin & Porphyrin |  |  |  | ✓ | ✓ |  | EoL? |

In summary, Bile acids and Steroids levels uniquely associated with EL were predictive of reduced mortality risk, supporting their potential role in EL, while the levels of selected Fatty Acids and Hemoglobin/Porphyrin metabolites uniquely associated with EL were predictive of increased mortality risk, suggesting they may reflect end-of-life processes.

### Metabolite Ratios Provide Mechanistic Insight Beyond Individual Metabolites

To develop insight into pathways that may underlie these findings, we then evaluated the association of a curated list of metabolite ratios with age and survival. Since metabolites are substrates and products of enzymatic reactions, the ratio of substrate to product may provide a more stable and informative measure than individual metabolites, as they can account for variations in sample concentration and better capture the balance of related metabolic processes. Metabolite ratios can thus serve as biomarkers for specific diseases or physiological conditions^39^.

We compiled a set of 267 ratios, with 69 that were manually curated based on our study’s findings, an additional set of 198 that were retrieved from literature corresponding to metabolite pairs sharing an enzyme or transporter *and* assayed in our dataset (Table S12a)^40^. Several of the manually curated ratios were selected based on the observed increased or decreased level with age of specific metabolites. For example, given the observed significant decrease with age of androgen- and progesterone-related steroids, as well as of glucocorticoids (cortisone), we investigated the ratio of cholesterol with metabolites in these classes. To investigate why dicarboxylic (DC) fatty acids (FA) increased with age (e.g., azelate, suberate, pimelate), we looked at their ratio with parent FAs (e.g., pelargonate, caprylate, heptanoate). Similarly, given the significantly higher level of bile acids (BA) in EL, we investigated whether glycine and taurine conjugation of BAs partially explained their EL-associated changes^41,42^. Finally, given the increase with age of N-acetylated amino acids (AA) (e.g., N-acetylaspartate, N-acetylleucine, N-acetylmethionine) we derived their ratios with parent AAs (e.g., aspartate, leucine, methionine).

Regression analysis of the log_2_-transformed ratios on age and covariates yielded a set of 33 and 24 ratios significantly (q ≤ 0.05) positively and negatively associated with age, respectively (Table S12b), with uridine / pseudouridine being the top ranked ratio negatively associated with age. Of note, tryptophan/kynurenine and aspartate/quinolinate, which contain metabolites within the *de novo NAD synthesis* pathway, were ranked as the 3^rd^ and 20^th^ ratios most significantly associated with age, respectively, with lower values associated with increased age, and both positively associated with uridine/pseudouridine, after adjusting for age and sex (ρ=0.534; q<1.0E-15 and ρ=0.507; q<1.0E-12, respectively). Additionally, the cholesterol/DHEA-S ratio was ranked 2^nd^ and positively associated with age, alongside multiple cholesterol/steroid ratios. Finally, the ergothioneine/TMAO and methionine/methionine sulfoxide ratios were ranked 16^th^ and 27^th^, respectively, and were both negatively associated with age (Table S12b).

Interestingly, many of the metabolites that were significantly associated with age and/or EL in NECS were also associated with the tryptophan/kynurenine, aspartate/quinolinate, ergothioneine/TMAO, methionine/methionine sulfoxide and cortisone/cortisol ratios, but opposite in direction (Table S12c), which suggests an underlying mechanistic role for these ratios on healthy aging and/or longevity.

### Metabolomic Clock Predicts Mortality Risk-Associated “Biological Age”

We evaluated the ability of metabolite levels to predict a subject’s *biological* age (BA), as distinct from the *chronological* age (CA) determined by a subject’s date of birth, with the difference between the predicted and the known CA (the *age deviation*, Δ) assumed to reflect variation in aging rates associated with mortality risk and age-related diseases^43^. Absolute and relative (i.e., bias-corrected)^44–47^ predictions by the fitted metabolomic clock are shown in Figure 5A. The clock included non-zero regression coefficients for 200 metabolites, with the top 25 identified metabolites ranked by permutation-based variable importance (VI) shown in Figure 5C (full list in Table S13). Survival analysis based on a Cox-PH model showed a significant association of the age deviation, Δ*(age)_corrected_*, with mortality risk: subjects predicted to be younger (older) than their chronological age showed a decreased (increased) mortality risk (e^β^ ≅ 1.085, p = 0.0196) (Figure 5D), thus confirming the ability of the metabolomic clock to capture salient information about a subject health status. Interestingly, while most of the clock’s highly predictive metabolites were also included in the age signature (previous section), some were not, with 6 of the top 25 top metabolites ranked by variable importance falling in this category. These included *1-methyl-5-imidazolelactate* (*β_m5i_* = −0.936, age q-value ≫ 0.05), the top ranked metabolite by VI and a member of the histidine metabolism pathway, followed by *3-hydroxyoctanoylcarnitine (1)* (VI rank = 3, *β_3hoc_* = −1.093, age q-value ≫ 0.05), *2’-o-methylcytidine* (VI rank = 8, *β_2’om_*= −0.163, age q-value ≫ 0.05), *s-carboxyethylcysteine* (VI rank = 17, *β_s-cec_* = 0.392, age q-value ≫ 0.05), and *taurine*, whose higher level was highly predictive of age deceleration (VI rank = 21, *β_taurine_* = −1.86, age q-value ≫ 0.05), and second only to *uridine* by negative size effect (*β_uridine_* = −2.025). While also associated with age, *citrate* was the 2^nd^ top ranked metabolite by variable importance, and was the top predictor of age acceleration by positive size effect (*β_citrate_* = 1.85)^48^.

**Figure 5.**
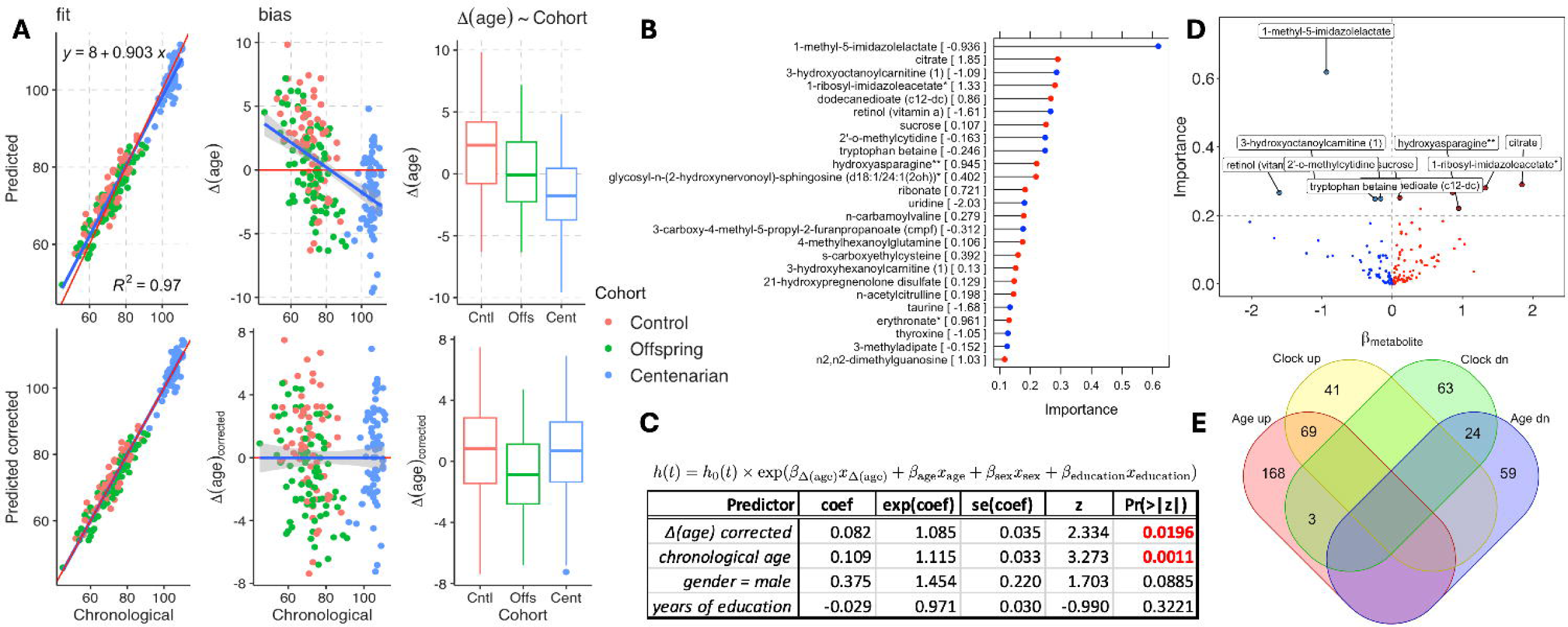
Metabolomics clock based on elastic net. **A**) before (top) and after (bottom) bias correction. **B**) Top 25 identified metabolites ranked by permutation-based variable importance (red/blue corresponding to positive/negative regression coefficients). **C**) Cox-PH-based survival analysis regressing on corrected delta-age (predicted – chronological). **D**) Volcano plot of elastic net metabolites’ beta estimates (x-axis) against importance values (y-axis). **E**) Overlap between significant aging signature metabolites and clock metabolites with non-zero beta estimates.

To evaluate the generalizability of the predictions, we repeated the analysis by 10-fold cross-validation (CV) (Figure S4). This approach yielded predictions with a smaller *R*^2^ (∼0.76) than the model trained on the complete dataset (∼0.97). However, the directions of the age deviations were mostly in agreement with those obtained from the model trained on the complete dataset (Figure S4C). Most importantly, the age deviations remained significantly associated with mortality risk (e^β^ ≅ 1.03, p-value < 0.025), which supports the conclusion that the age deviations’ relative *trends*, rather than their absolute magnitude are predictive of health status.

## DISCUSSION

In this study, we performed metabolomic profiling of a cohort significantly enriched for exceptionally long-lived (EL) subjects. While previous studies included centenarians, none included ages as advanced as those represented in NECS, with median, 75^th^-percentile, and maximum ages at the time of blood draw of 75y, 103y, and 110y, respectively, reaching 88y, 106y, and 115y at last follow-up.

Methodologically, evaluation of metabolites as classes increased the statistical power of our analyses, which led to the discovery of associations that would have been otherwise overlooked. Additionally, integration of our EL-enriched cohort with results from independent non-EL studies, and triangulation with survival, enabled us to zoom-in on what was likely *unique* to aging *vs.* EL, and to focus on the identification of markers of either. The results of our analyses confirm and significantly expand upon previous findings, and we discuss them below.

*<u>Aging-Associated Metabolites Reflect Known Hallmarks</u>*. Metabolites that increase with age are enriched in pathways related to key aging hallmarks such as oxidative stress, inflammation, mitochondrial dysfunction, and renal decline, including urea cycle intermediates (e.g., citrulline, proline), betaine derivatives linked to one-carbon metabolism and epigenetic aging, and markers of protein glycation and oxidative damage ^14,26–28,49^. Additionally, tyrosine metabolites and dicarboxylic acids associated with mitochondrial and renal dysfunction also accumulate with age, correlating with findings in centenarians and suggesting their relevance to longevity ^29–37,50^. Conversely, several lipid classes, including long chain polyunsaturated fatty acids (PUFAs) and certain acyl choline and monounsaturated fatty acids (MUFAs), decline with age, though patterns vary between extremely long-lived (EL) and non-EL individuals, highlighting complex roles of lipid metabolism in aging.^34,36,51–57^

*<u>EL-only markers</u>*. Metabolites uniquely associated with EL revealed distinct biochemical features not observed in typical aging trajectories. Most notably, elevated levels of *primary* and *secondary bile acids* in EL participants – particularly those containing lithocholic acid (LCA) – were associated with improved survival. These findings are consistent with prior work identifying bile acid biosynthesis enrichment in centenarians^20,21^, and extend those results by showing a potential survival benefit. They also lend human relevance to recent studies in long-lived model organisms, which showed that the level of LCA rises with calorie restriction and prolongs lifespan^58,59^, and that circulating levels of bile acids are elevated in dwarf mice^60^, a model of extended lifespan^61^.

By contrast, EL subjects exhibited reduced levels of *long-chain PUFAs*, *MUFAs*, and *acylcholines* relative to younger cohorts. Some of these changes (e.g., decreased acylcholines) were paradoxically linked to decreased survival, potentially marking physiological frailty or end-of-life changes.

Several *heme-related metabolites*, including biliverdin and bilirubin isomers, were significantly reduced in EL, suggesting possible disruption of the redox cycle and antioxidant defenses, perhaps due to biliverdin reductase dysfunction^62,63^. This observation contrasts with some prior reports of increased bilirubin levels with age^64^ and raises intriguing questions about the balance between antioxidant protection and oxidative stress regulation in the extremely old.

*<u>Age-only markers</u>*. Metabolites uniquely associated with chronological age – but not with EL – highlight potential biomarkers of typical aging trajectories rather than healthy or exceptional aging. Among these, *acylcarnitines* from several lipid subclasses showed consistent increases with age, consistent with prior studies^27,65–67^, with elevated levels associated with mitochondrial dysfunction^68–70^. However, this increase was not maintained in EL individuals, nor were they significantly associated with survival, suggesting that while they may track chronological aging, they do not necessarily reflect healthy aging. Previous findings have suggested an inverse relationship between acylcarnitines and biological age^56^, supporting the notion that lower levels may be beneficial.

Interestingly, several *steroid classes*, primarily progestins, and selected androgens and pregnenolone derivatives, declined with age but remained stable in EL individuals. Higher levels of these steroids were associated with lower mortality risk, suggesting a possible protective role. Positive associations between these steroids and the tryptophan/kynurenine ratio further support their relevance to immune and metabolic regulation, as a lower level of this ratio is a recognized marker of systemic inflammation and immune activation^71,72^. Similarly, positive associations of the tryptophan/kynurenine ratio with metabolites from the hemoglobin and steroid metabolism pathways, and opposite in trend with their age-related association, were also observed. The importance of tryptophan/kynurenine and other metabolite ratios that were significantly associated with age, EL, and/or survival in our study is discussed below.

### A potential role for NAD (and/or inflammation) and gut bacterial metabolism in healthy aging

Multiple metabolites from the *de novo* NAD synthesis pathway were significantly associated with age and EL, including positive associations for kynurenine and quinolinate. In contrast, tryptophan was negatively associated with both age and, marginally, with EL (q = 0.07). A declining tryptophan/kynurenine ratio with age is consistent with previous reports linking reduced levels of this ratio to elevated inflammation and/or reduced NAD synthesis efficiency^71,72^, which suggests age-associated increased inflammation and/or low NAD in the NECS cohort. Given NAD’s central role in aging and age-related disease prevention^73^, we further investigated age-associated metabolites that were also associated with the tryptophan/kynurenine ratio (while controlling for age). Metabolites from a variety of pathways that increased during aging and in EL were negatively associated with the tryptophan/kynurenine ratio (Table S12c), which suggests a potential role for sufficient NAD and/or low inflammation on healthy longevity. Integration of the metabolite ratio results with study findings suggested a potential feedback loop linking NAD metabolism, inflammation, and gut microbiome activity (Figure 6).

**Figure 6.**
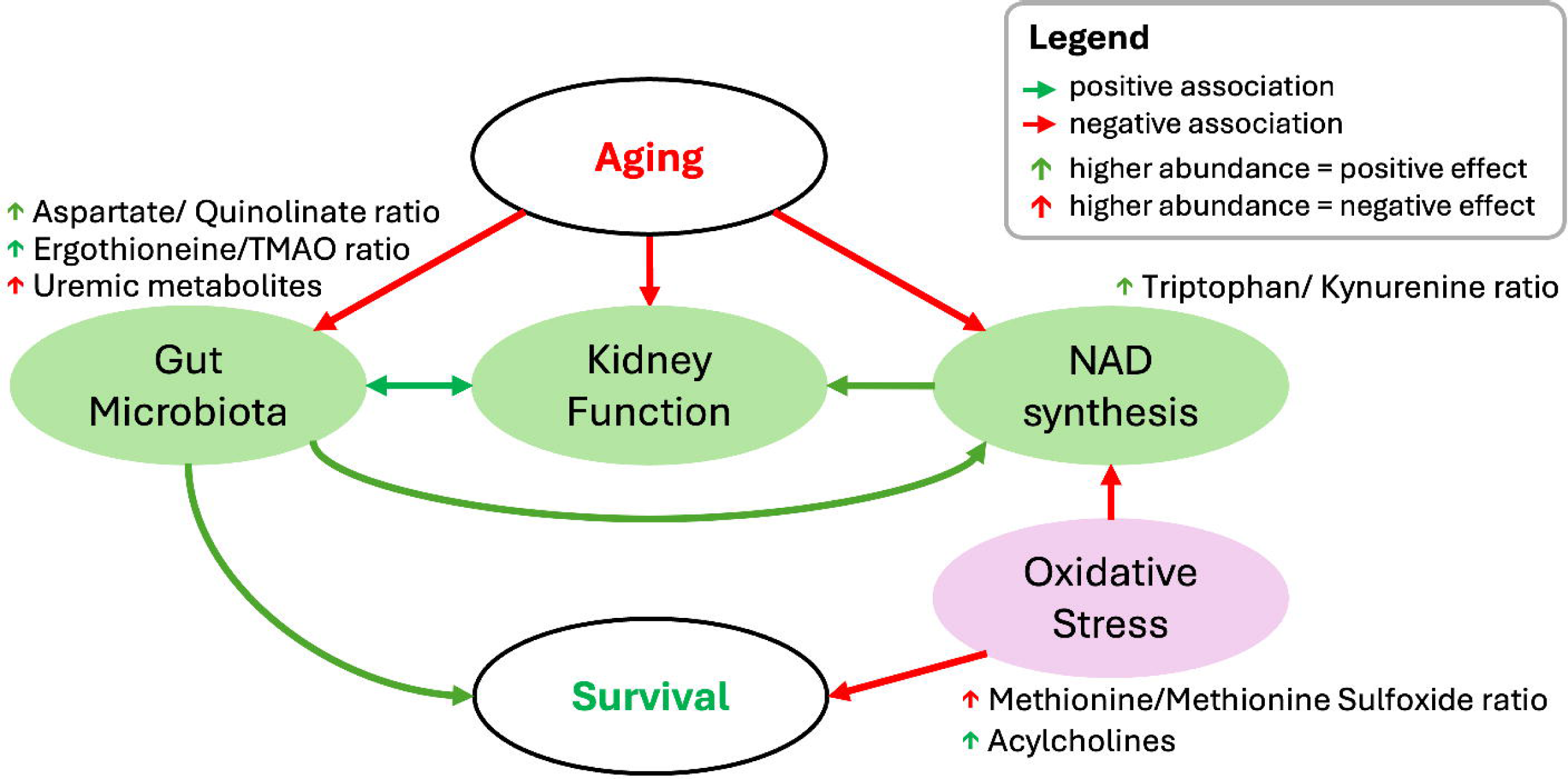
A feedback loop linking healthy aging to NAD metabolism, gut microbiome activity, renal function, and oxidative stress. Markers of increased NAD synthesis and/or reduced inflammation (tryptophan/kynurenine), a healthy gut bacterial metabolism (aspartate/quinolinate, ergothioneine/TMAO), and reduced oxidative stress (methionine/methionine sulfoxide) are negatively associated with many metabolites strongly associated with age or with worse survival.

*Uremic metabolites increasing with age are negatively associated with a biomarker of reduced inflammation/ increased NAD synthesis*. Several metabolites that increased with age and EL – including phenol sulfate, p-cresol glucuronide, multiple sugar acids, and various N-acetyl amino acids – were negatively associated with the tryptophan/kynurenine ratio. Higher levels of sugar acids were also associated with shorter survival (q = 0.08, Table S11). These metabolites are known uremic toxins, or are associated with a reduced eGFR^50,74–77^, implicating a relatively high tryptophan/kynurenine ratio as a potential marker of healthy renal function during aging. This is supported by prior studies reporting an inverse association between the ratio and incident chronic kidney disease^78^.

*Quinolinate accumulation with age is associated with biomarkers of gut bacterial metabolism and increased inflammation/reduced NAD synthesis*. Quinolinate, a downstream product in the *de novo* NAD synthesis pathway, is also implicated in neuroinflammatory conditions such as Alzheimer’s disease^79^. Its increased levels with age and in EL may reflect impaired NAD synthesis, as indicated by the concurrent decline in the tryptophan/kynurenine ratio. In contrast, here we tested the novel hypothesis that increased gut bacterial metabolism contributes to quinolinate accumulation with age. Aspartate is converted into quinolinate via L-aspartate oxidase, an enzyme that is specific to bacteria^80^, including *Escherichia coli*, whose gut levels increase with age^81^. Accordingly, a lower aspartate/quinolinate ratio is a marker of increased bacterial aspartate degradation into quinolinate, and conversely, a higher ratio is indicative of less bacterial conversion. Of note, aspartate/quinolinate was positively associated with tryptophan/kynurenine (while controlling for age) in our data. Moreover, many of the metabolites that were associated with age and EL, but were negatively associated with tryptophan/kynurenine, were also negatively associated with aspartate/quinolinate, which further supports a link between gut bacterial metabolism and NAD, inflammation, aging, and/or EL (Table S12b,c)^82–85^.

*Acylcholines decline during aging, and higher levels are associated with biomarkers of gut bacterial metabolism and reduced oxidative stress*. Acylcholines declined with age and in EL, and higher levels were associated with improved survival. Higher acylcholine levels were positively associated with aspartate/quinolinlate – a biomarker of gut bacterial metabolism – and with the methionine/methionine sulfoxide ratio – a biomarker of lower oxidative stress (i.e., higher ratio values reflect reduced oxidative stress). Aspartate/quinolinate’s positive association with acylcholines suggests that reduced microbial degradation of aspartate to quinolinate may support acylcholine-related longevity mechanisms. Supporting their protective role, long-chain acylcholines are elevated in HIV-1 patients that are able to suppress viremia without antiretroviral therapy (which suggests a role for acylcholines on immune defense) and are linked to lower type II diabetes risk^86^, highlighting their potential importance in age-related phenotypes.

*Ergothioneine’s decline with aging is negatively associated with biomarkers of gut bacterial metabolism and reduced inflammation/increased NAD synthesis.* Ergothioneine is a metabolite linked to lifespan extension in animal models and has been proposed as a ‘longevity vitamin.’ ^87–89^ In the present study, ergothioneine levels decreased with age and extreme longevity, confirming our previous study^22^. It is abundant in mushrooms but can be degraded by gut bacteria into trimethylamine (TMA), which the liver converts to trimethylamine N-oxide (TMAO)^90^. The ergothioneine/TMAO ratio reflects gut bacterial activity, with a higher ratio indicating lower breakdown by bacteria. This ratio was negatively associated with harmful uremic metabolites that increase with age, suggesting gut bacteria influence kidney function and longevity^91^. Moreover, a higher ergothioneine/TMAO ratio correlated positively with steroids and hemoglobin-related metabolites, which decline with age but support survival. It also correlated with the tryptophan/kynurenine ratio, a marker of NAD metabolism and inflammation, raising questions about whether gut bacteria influence metabolic health or vice versa.

Taken together, these results show that markers of decreased gut bacterial activity (higher aspartate/quinolinate and ergothioneine/TMAO ratios), increased NAD synthesis and/or reduced inflammation (higher tryptophan/ kynurenine and cortisone/cortisol ratios), and reduced oxidative stress (higher methionine/methionine sulfoxide ratio) are negatively associated with many metabolites strongly associated with age or with worse survival, which suggests an underlying role for these pathways in aging and, potentially, longevity.

### Metabolomic Clock Reveals Biomarkers of Biological Aging

The metabolomic clock that we developed offers a promising tool for estimating biological age and identifying individuals with accelerated or decelerated aging. The significant association between corrected age deviation and mortality supports its utility as a marker of healthy aging. Notably, several metabolites with high predictive value were not significantly associated with chronological age. The presence of predictive metabolites not captured in traditional age-association analyses suggests that the clock can reveal aging-relevant biology that may be missed when focusing solely on cross-sectional univariate age correlations. Notably, taurine, which was highly predictive of age deceleration, aligns with recent experimental studies linking taurine supplementation to increased lifespan in animal models, raising the possibility of translational relevance^42,92^, although a recent observational study found that circulating taurine concentrations increased or remained unchanged with age in human cohorts as well as in nonhuman primates and mice when measured longitudinally^93^. Similarly, citrate, the strongest positive predictor of age acceleration, has been implicated in mitochondrial and inflammatory pathways associated with aging, and part of a six-metabolite score highly predictive of mortality^48,94^. Taken together, these findings highlight the potential of metabolomic clocks not only to quantify biological age, but also to prioritize specific metabolic pathways and molecules for therapeutic targeting.

### Study Limitations

Despite the comprehensive nature of this untargeted metabolomic analysis and integration across multiple cohorts, some limitations should be acknowledged. First, the cross-sectional design limits our ability to infer causality or temporal dynamics of metabolite changes during aging and extreme longevity. Longitudinal metabolomic profiling will be critical to differentiate stable longevity signatures from end-of-life metabolic shifts. Second, although the NECS cohort is uniquely enriched for exceptionally long-lived individuals, the sample size of centenarians remains relatively modest, which may reduce power to detect rarer metabolic changes and limits generalizability to broader aging populations. Third, differences in metabolomic platforms, batch effects, and coverage across integrated datasets posed analytical challenges and could contribute to incomplete replication of findings. Additionally, the influence of unmeasured lifestyle, dietary, and medication factors may confound metabolite associations with longevity and survival. Finally, while our metabolomic clock shows promise in predicting biological age and mortality risk, further validation in independent, ethnically diverse cohorts with long follow-up is needed to confirm its robustness and translational utility. Future studies addressing these limitations will help clarify the mechanistic roles of identified metabolites in healthy aging and inform targeted interventions.

## MATERIALS & METHODS

### Human Cohorts, Metabolomic, and Phenotypic Data

*<u>New England Centenarian Study (NECS)</u>*. ***Cohort description.*** Metabolomic profiles of 213 NECS participants comprising centenarians (n=80), centenarians’ offspring (n=70) and unrelated controls^17^ (n=63) were included for analysis. Centenarians were defined as those subjects achieving ages above the 99^th^ percentile of their age cohort, with all but one (male) reaching > 100-year of age at last follow-up. We selected participants who were at least one year away from major clinical events, free of major medications, and survived at least one year after the blood draw. Trained phlebotomists collected blood samples from NECS participants at enrollment and transferred the samples to the Boston University Molecular Genetic Core biobank for DNA, serum, and plasma extractions. Samples were maintained at −80°C. ***Metabolomic data.*** Metabolon, Inc. (North Carolina, USA) generated serum metabolomic profiles, which included 1,495 chemicals (1,213 and 282 compounds of known and unknown identity, respectively). A detailed description of sample preparation and liquid chromatography-tandem/mass spectrometry (LC/MS) was previously described^23^. The NECS protocol was approved by the Boston Medical Center Institutional Review Board. Participants provided written informed consent at enrollment.

*<u>Arivale</u>. **Cohort description.*** The Arivale cohort includes adults (age range 18-90+) who self-enrolled in a consumer-facing wellness program (Arivale, Inc. 2015-2019)^95^. The participants were required to be over the age of 18 and not pregnant, with no additional screening of participants. Certified phlebotomists collected blood samples from fasting participants. We included a subset of 634 participants from the Arivale cohort with plasma metabolomic and other covariate data. ***Metabolomic data.*** Metabolon, Inc. (North Carolina) generated plasma metabolomic profiles using ultra-high-performance liquid chromatography–tandem mass spectrometry. Sample handling, quality control, data extraction, biochemical identification, data curation, quantification, data normalizations and batch correction have been previously described^24,96^. After name curation, 615 metabolites were matched to metabolites in the NECS dataset. The current study was conducted with de-identified data of the participants who had consented to the sharing of their deidentified data. Procedures were approved by the Western Institutional Review Board.

*<u>Long Life Family Study (LLFS)</u>. **Cohort description***. The LLFS is a family-based, longitudinal study of healthy aging and longevity that enrolled 4,953 family members from 539 families selected for familial longevity at three American field centers (Boston, Pittsburgh and New York), and a Danish field center between 2006 and 2009^97,98^. Study participants have been comprehensively phenotyped using a combination of three in-person visits, questionnaire-based assessments, and annual follow-ups. ***Metabolomic data.*** A total of 408 metabolites (188 lipid and 220 polar metabolites) were profiled in the plasma of 2,764 LLFS participants collected at enrollment. Of these, 163 metabolites were matched to the NECS metabolites based on RefMet standardized names^99^. The LLFS experimental workflows have previously been described^23,100^.

*<u>Baltimore Longitudinal Study of Aging (BLSA)</u>*. ***Cohort description.*** The BLSA is a longitudinal study of community-dwelling adults that continuously enrolls participants free of cognitive impairment, functional limitations, chronic diseases, and cancer for the previous 10 years^25^. Details of blood collections have been previously described^101,102^. We selected a subset of 1,135 BLSA participants with plasma metabolomic data, including a subset of 926 participants aged ≥50 years with follow-up. ***Metabolomic data.*** Metabolite concentrations were measured via the Biocrates p500 kit with LC-MS/MS. (Biocrates Life Sciences AG, Innsbruck, Austria), and following the manufacturer’s protocol for a 5500 QTrap (Sciex, Framingham, MA, USA). Ninety-nine metabolites were in common with the LLFS using RefMet standardized names^99^. The BLSA protocol was approved by the Institutional Review Board of the National Institutes of Health. Participants provided written informed consent at each BLSA visit.

*<u>Xu et al cohort</u>*. ***Cohort description.*** Xu *et al.* described multi-omic profiles for 116 nonagenarians and centenarians, and 232 of their offspring from Han Chinese longevous families^21^. ***Metabolomic data.*** 821 lipid and polar metabolites were generated using non-targeted metabolomic analysis, and 432 metabolites were matched to metabolites in the NECS dataset. Study details are described in the manuscript and individual level data are available in Table S6 of their supplementary material^21^.

#### Datasets Harmonization

The datasets analyzed were generated based on different platforms and used different identifiers for the metabolites measured. To map metabolites across datasets, we used different approaches depending on the datasets to be matched. For example, when comparing NECS with Arivale and with Xu *et al*., we used the Metabolon-provided CHEMICAL_NAME’s, which yielded an overlap of 615 and 385 metabolites, respectively. When comparing NECS with LLFS and BLSA, we mapped all metabolites to RefMet identifiers^99^, which yielded an overlap of 163 and 188 metabolites, respectively. It should be emphasized that the integration of the NECS cohorts with the other datasets was performed at the level of the summary statistics (i.e., beta estimates and adjusted p-values from the age and EL regression analyses).

### Metabolite Sets (MSets) Definition

Annotation of the metabolite signatures was performed using metabolite sets (MSets) we defined based on the Metabolon-provided annotation, as well as the RefMet annotation^99^. Metabolon annotation includes a two-level hierarchy grouping metabolites into *SUPER_PATHWAY* and *SUB_PATHWAY* categories, with the latter nested in the former. Based on the annotated metabolites in our datasets, and using the *SUB_PATHWAY* categorization, we created a compendium of 92 (mutually exclusive) MSets containing between 3 and 91 metabolites. Similarly, RefMet provides a three-level hierarchy grouping metabolites into *super_class*, *main_class*, and *sub_class*, and we used the latter to create a compendium of 71 (mutually exclusive) MSets containing between 3 and 148 metabolites. These two compendia were used throughout the study for signature and (WGCNA) module enrichment analysis. They are also available as “hierarchical rgsets” (see hypeR documentation) in the GitHub associated with this study. Sample-specific Metabolon and RefMet MSet scores (Figure S3) were computed using GSVA^38^ using the log_2_-transformed metabolite levels as input.

### Metabolite Module Detection

For the detection of modules of highly correlated metabolites, we applied WGCNA^103^ considering only positive correlations using the *bicor* function. Average linkage was used as the hierarchical clustering agglomeration rule. Modules were constrained to have a minimum size of 10, and modules “too close as measured by the correlation of their eigengenes” were merged (using the *merge = TRUE* option). To control for confounders, WGCNA was applied to the matrix of residuals after regressing on age, serum’s length of storage, sex, years of education, and batch. Sample-specific module scores (Figure 2) were computed using GSVA^38^ based on the log_2_-transformed metabolite levels.

### Statistical Analysis

#### Age and EL regression

For the identification of age-associated metabolites, we fitted a linear least squares (LLS) model, regressing the log2-transformed metabolite level on age while controlling for *sex*, storage duration of *serum*, years of *education*, and *batch*.

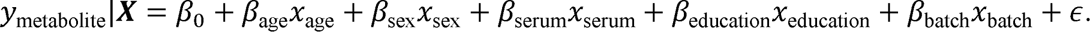

Metabolites with an FDR-adjusted *β*_age_ coefficient q-value ≤ 0.05 were selected, with a *β*_age_ >0 indicating metabolites positively associated with age (i.e., higher levels associated with older age) and vice versa.

Similarly, for the identification of EL-associated metabolites, we regressed on a categorical 3-valued variable (control, offspring, EL, with control as the reference) while controlling for the same set of covariates.

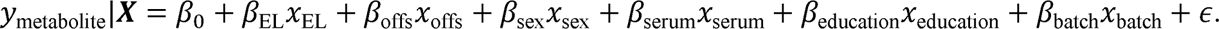

Metabolites with an FDR-adjusted *β*_EL_ coefficient q-value ≤ 0.05 were selected, with a *β*_EL_ >0 indicating metabolites with higher levels in the EL group and vice versa.

#### Survival analysis

To identify metabolites associated with mortality risk, we fitted a Cox’s Proportional Hazards (CoxPH) model regressing right-censored length of survival on metabolite level, while controlling for the same covariates as in the LLS models.

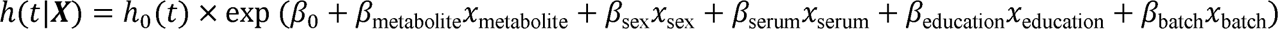

Testing of the proportional hazard assumption was performed using the *cox.zph* function in R. When performing rank-based MSet enrichment analysis against survival (see below), metabolites were ranked by their *β*_metabolite_ coefficient. Similar models were fitted when evaluating the metabolomics clock, with Δ*(age)* or Δ*(age)_corrected_* in place of the metabolite level in the above CoxPH equation.

#### Metabolite Sets (*MSets*) Enrichment Analysis

When performing MSet enrichment based on over-representation analysis (ORA), we used the hypeR package^104^ with test = “hyper”, setting the background to 1,213, which is the number of metabolites with known identity in the NECS dataset. When performing rank-based Kolmogorov-Smirnov MSet enrichment, we used the hypeR package with test = “ks” and using the entire list of 1,053 metabolites with ≤ 20% of missing values analyzed and ranked by their test statistic (e.g., the CoxPH coefficients in the survival analysis).

#### Definition of age-only and EL-only signatures

To identify age-only markers, we compared the set of EL-associated metabolites in NECS with age markers from datasets that did not include EL subjects (non-EL studies: Arivale and BLSA), and to the age markers obtained by performing regression analysis on the subset of NECS subjects not in the EL group, i.e., the set of 60 controls and 73 offspring. We then defined the *Age-only signature* as the set of metabolites that were significantly associated with age in either Arivale or BLSA (q ≤ 0.05), that had age estimates in NECS (excluding EL subjects) in the same direction (i.e., positive in NECS if positive in Arivale or BLSA, and vice versa), and had non-significant EL estimates (q ≥ 0.25) in NECS.

Similarly, the *EL-only signature* was defined as the set of metabolites that were significantly associated with EL (q ≤ 0.05) in NECS but not significantly associated with age (q > 0.25) in all non-EL studies in which they were assayed. Since not all metabolites assayed in NECS were measured in either Arivale or BLSA, we further restricted the EL-only signature to metabolites measured both in NECS and in either Arivale or BLSA.

#### Metabolomics Clocks

Metabolomics clocks based on an elastic net (EN) model were fitted using the *glm.net* function in R. Ten-fold cross-validation (CV) was used to select the α and λ coefficients maximizing the RMSE from a grid of 25 predefined alpha and lambda values’ combinations. The set of 1,053 metabolites with ≤ 20% of missing values were used as predictors, with the EN (non-zero) weights determining which metabolites to include in the final clock. Predictions by the model showed a moderate bias, with (chronologically) younger subjects predicted to be older and older subjects predicted to be younger (Figure 5A). The observed bias is common to most omics clocks^45–47^, and was corrected by a simple linear transformation^44^. Permutation-based variable importance for each of the metabolites with non-zero coefficients was estimated using the *vip* R package. Model fitting and prediction was also repeated based on a nested two-level 10-fold CV, with the outer CV loop used for age prediction, and the inner CV loop used for the selection of the optimal EN hyper-parameters (α and λ) for each fold (Figure S4).

---

**Figure S1.**
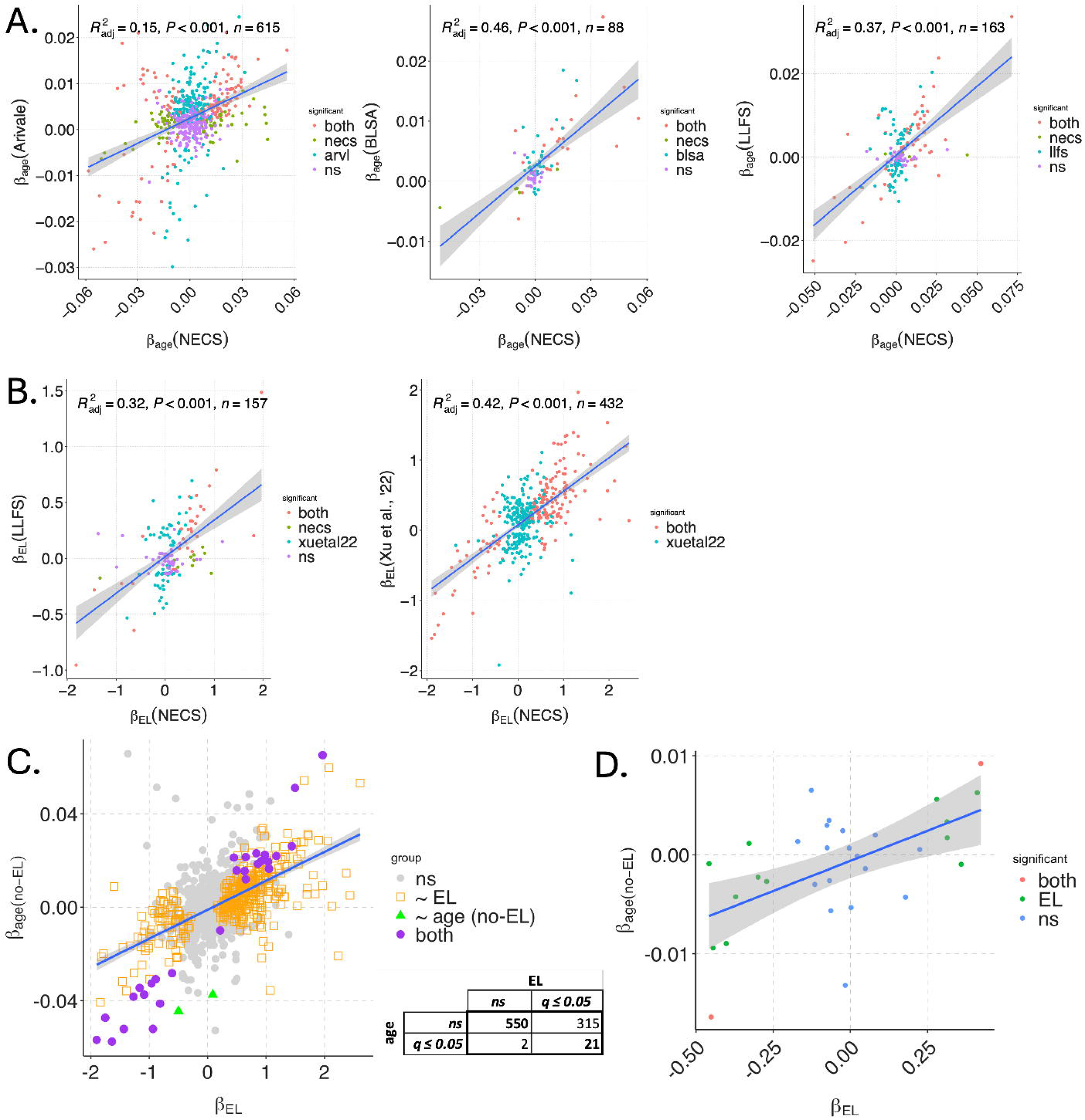
Comparison of beta estimates across datasets for (A) age and (B) EL regression models.

**Figure S2:**
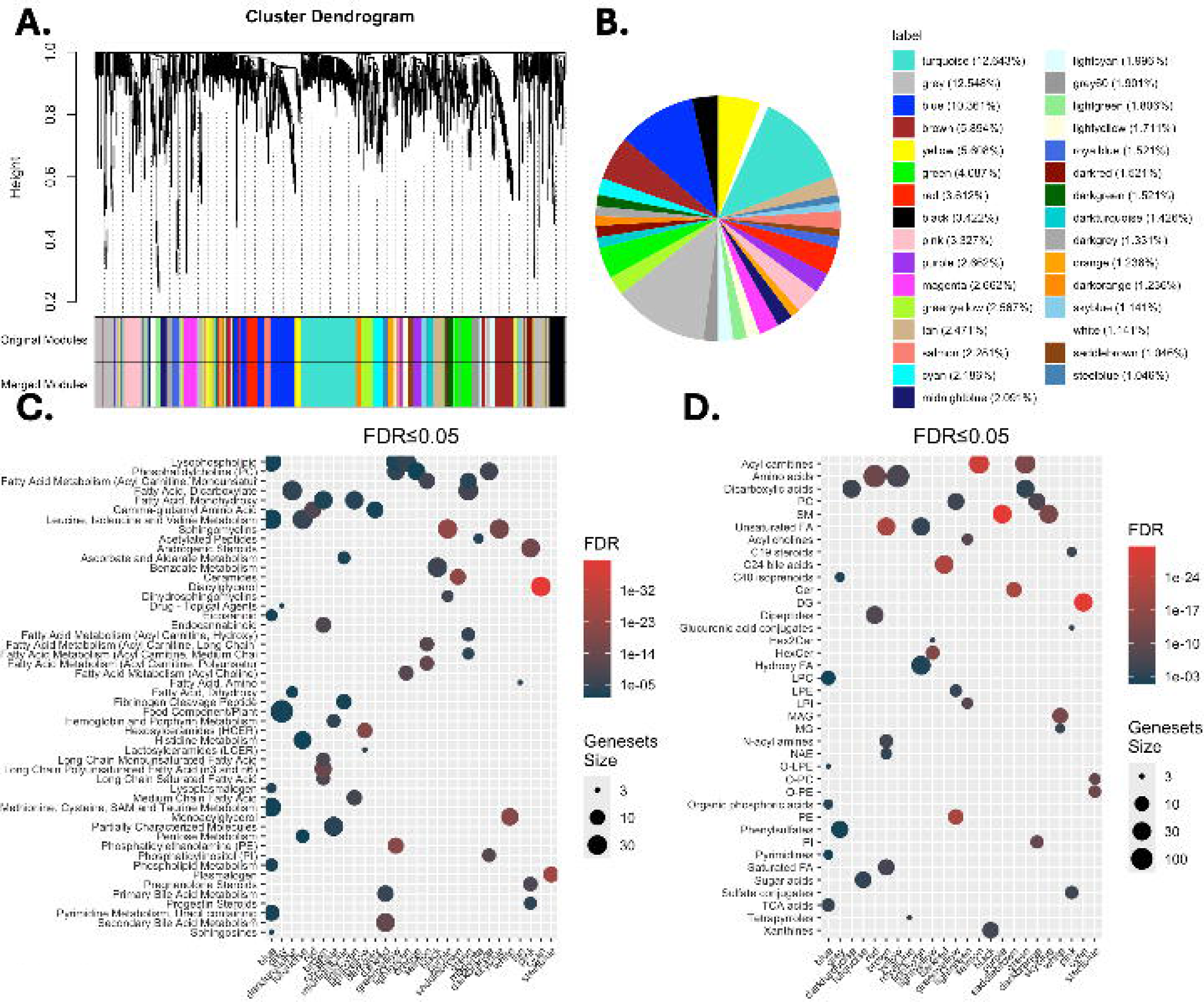
WGCNA-based module detection and annotation. **A)** Cluster dendrogram based on metabolites’ (positive) correlation, and identification of modules by dynamic tree cutting, followed by merging of modules. **B)** Modules’ relative size. **C-D)** Modules’ ORA-based enrichment based on Metabolon and RefMet MSets, respectively.

**Figure S3.**
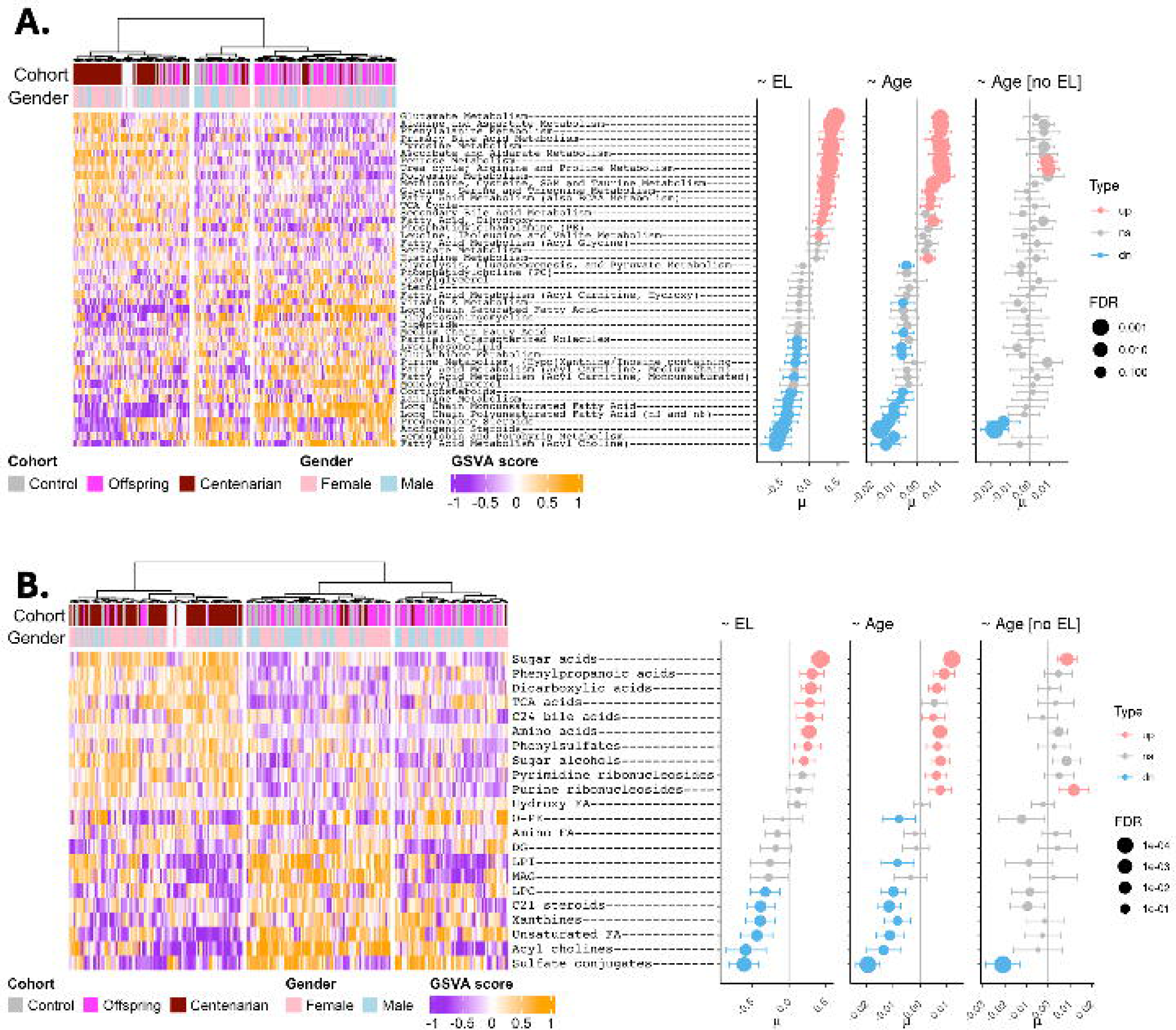
Module-based heatmaps (left) and regression analyses (right) on EL status and age (with and w/o EL subjects included), with sample-level module scores computed using GSVA. **A)** Modules (y-axis) represent Metabolon (SUB_PATHWAY) MSets. **B)** Modules (y-axis) represent RefMet (sub-class) MSets.

**Fig S4:**
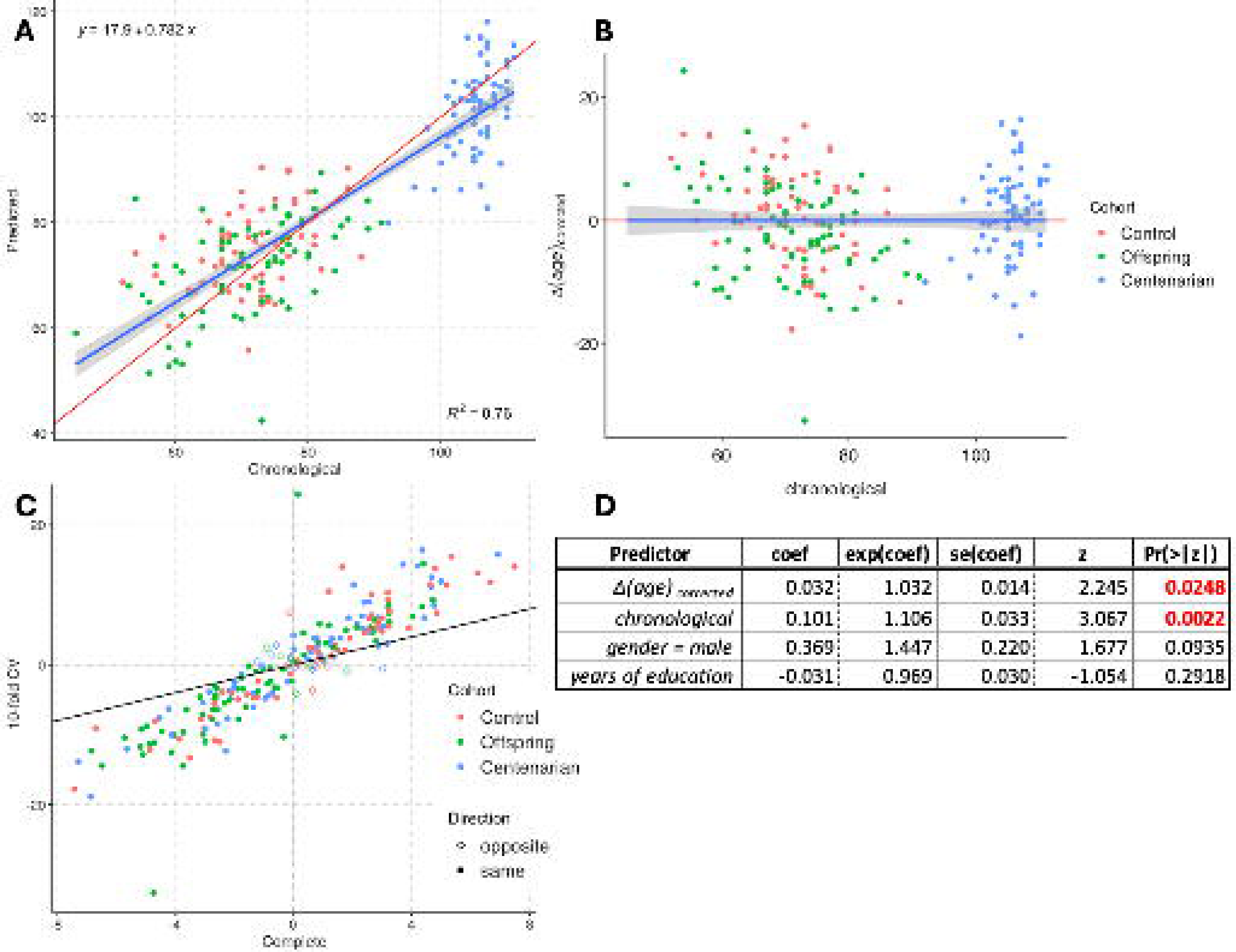
Metabolomics clock 10-fold cross-validation. **A**) Prediction and fit. **B**) Δ(age)_corrected_ of the 10-fold CV predictions. **C**) Comparison of Δ(age)_corrected_ of the predictions based on the models fitted on the Complete dataset and by 10-fold CV. **D**) Cox-PH-based survival analysis regressing on Δ(age)_corrected_ from the 10-fold CV model.

## SUPPLEMENTAL INFORMATION

**Table S1.** Annotated list of profiled metabolites.

**Table S2**. Summary statistics of the datasets included in the study. The column “# Overlap” indicates the number of metabolites in that dataset overlapping with the NECS metabolites.

**Table S3**. Age, Age (no-EL), EL, and survival metabolites’ signatures derived in NECS, and their replication across the other datasets (Arivale, BLSA, LLFS, Xu et al.).

**Table S4**. Over-representation analysis (ORA) of the metabolites’ signatures based on Metabolon’s *SUPER_PATHWAY*/*SUB_PATHWAY* MSets.

**Table S5**. Over-representation analysis (ORA) of the metabolites’ signatures based on RefMet’s *super_class*/*main_class*/*sub_class* MSets.

**Table S6**. WGCNA module composition.

**Table S7**. Age, age (no-EL), and EL signatures based on WGCNA-based module scores.

**Table S8**. Age, age (no-EL), and EL signatures based on Metabolon-based MSets scores.

**Table S9**. Age, age (no-EL), and EL signatures based on RefMet-based MSets scores.

**Table S10**. Rank-based enrichment analysis of the survival signature based on Metabolon MSets.

**Table S11**. Rank-based enrichment analysis of the survival signature based on RefMet MSets.

**Table S12**. Metabolite Ratios (a), their association with age, EL, and survival (b), and their correlation with other ratios and metabolites.

**Table S13**. Metabolomics clock’s metabolites, their variable importance, and their beta estimates.

## ACKNOWLEDGMENTS

This work was supported by the National Institutes of Health, NIA cooperative agreements U19-AG023122, U19AG063893, UH2AG064704, and NIH grant R01-AG061844.

## AUTHOR CONTRIBUTIONS

Conceptualization, S.M. and P.S.; data curation, S.M., P.S., Z. S., D.E., Q.T., N.R., L.F.; data analysis, S.M., Z.H., Z.S., P.S.; software, S.M. and Z.H.; results interpretation, S.M, M.S.L., L.F., P.S.; manuscript writing, original draft, S.M and M.S.L.; writing – review & editing, all authors.

